# miRCoop: Identifying Cooperating miRNAs via Kernel Based Interaction Tests

**DOI:** 10.1101/769307

**Authors:** Gulden Olgun, Oznur Tastan

## Abstract

Although miRNAs can cause widespread changes in expression programs, single miRNAs typically induce mild repression on their targets. Cooperativity is reported as one strategy to overcome this constraint. Expanding the catalog of synergistic miRNAs is critical for understanding gene regulation and for developing miRNA-based therapeutics. In this study, we develop miRCoop to identify synergistic miRNA pairs that have weak or no repression on the target mRNA, but when bound together, induce strong repression. miRCoop uses kernel-based interaction tests together with miRNA and mRNA target information. We apply our approach to kidney tumor patient data and identify 66 putative triplets. For 64 of these triplets, there is at least one common transcription factor that potentially regulates all participating RNAs of the triplet, supporting a functional association among them. Furthermore, we find that triplets are enriched for certain biological processes that are relevant to kidney cancer. Some of the synergistic miRNAs are very closely encoded in the genome, hinting a functional association among them. We believe miRCoop can aid our understanding of the complex regulatory interactions in different health and disease states of the cell and can help in designing miRNA-based therapies. Matlab code for the methodology is provided in https://github.com/guldenolgun/miRCoop.

## 1 Introduction

MicroRNAs (miRNAs) are small non-coding RNAs that play extensive regulatory roles across different biological processes and pathologies such as cancer ([18, 44]). By binding the complementary sequence of the target messenger RNAs (mRNAs), miRNAs repress their target mRNA gene expression ([5]). Despite the widespread changes they have on the protein levels, single miRNAs typically induce modest repression on their targets ([49, 4]). Cooperation among miRNAs is one strategy to boost their activity. This cooperativity can be realized in the form of multiple miRNAs collectively co-regulating a cohort of genes that are part of a functional module, or multiple miRNAs can jointly co-target an individual gene to achieve cooperative control of a target gene (reviewed in [9]). In this work, we focus on the latter case.

There is a substantial amount of experimental evidence on the cooperativity of miRNAs where a stronger degree of repression of a single gene is achieved with multiple miRNAs than is possible by the individual miRNAs. Enright et al. [16] are the first to show the cooperativity of lin–4 and let–7 in Drosophila experimentally. Krek et al. [32] find that in a murine pancreatic cell line that miR–375, miR–124, and let–7b cooperate to repress the MTPN gene. Similarly, Lai et al. [33] reports synergistic interaction of miR–93 and miR– 572 in repressing the p21 gene in human SK–Mel– 147 melanoma cells. Yang et al. [61] shows that miR–17–5p and miR–17–3p can synergistically induce prostate tumor growth and invasion by repressing the same target TIMP3. Others also show similar lines of evidence on the synergistic interactions of miRNAs to regulate a mutual target ([20, 57, 55, 54, 34]).

Despite the accumulating evidence, our understanding of synergistic miRNAs that regulate a single gene is still rudimentary. Determining which pairs of miRNAs synergize and which mRNA is the target of this cooperativity is critical for understanding how miRNAs function and how they regulate various gene expression programs. Furthermore, synergism can be exploited in designing miRNA based therapeutics as pointed in a recent survey by Lai et al. [34]. The single miRNA treatments require high amounts of miRNAs which can provoke off-target effects while therapeutics based on synergistic miRNAs have the potential to overcome these unintended effects by enabling lower levels of each of the miRNAs to be administered ([34]).

In this study, we develop miRCoop for detecting synergistic miRNAs and their mutual targets using the expression data and miRNA target information. We focus on a specific type of synergistic relationship: the miRNA pairs which have weak or no repression on the target mRNA, but when together strongly repress its expression. Please note that this does not cover all the synergistic relationships, i.e., the case where miRNAs have moderate expression themselves can also be in a synergistic relationship. Although there are methods that address the cooperativity of miRNAs - discussed below - there is no work that specifically addresses this specific type of synergistic relationship. We focus on this particular case because this set of miRNAs can uncover more novel RNAs that are important for a disease or condition which cannot be captured by merely checking the miRNAs individual effect, and they are important for miRNA-based therapies.

Several related computational methods exist in the literature on miRNA cooperativity. However, the majority of these tools inspect the cooperativity of miRNAs in regulating *a set of genes* which are part of a functional module ([50, 64, 3, 8, 2, 59, 36, 21, 13, 40, 14]) in contrast to the cooperativity for regulating *a single gene*. Hon et al. [25] is one of the early work that analyzes the combinatorial interactions of miRNAs and their targets. Although the study includes an analysis of the miRNAs that mutually target a single gene, this work does not aim at finding miRNAs that bind their target concurrently in a synergistic manner. The target relationships are established solely based on the predicted miRNA-mRNA interactions without expression data.

TriplexRNA aims at detecting synergistic miRNAs that bind to a mutual target gene [47]. TriplexRNA assesses the synergism of the pairs of miRNAs in the human genome using miRNA target predictions, structural predictions, molecular dynamics simulations, and mathematical modeling methods. The main limitation of this method is that it does not incorporate the gene expression profiles. The gene regulation changes drastically in different conditions; thus, the identified miRNAs are putative triplets and are not context-dependent. CancerNet [40] makes use of the expression data but not for testing the statistical interactions of the triplets. The miRNA-mRNA expression profiles in each cancer are used to remove the targets that are not expressed from the protein-protein interaction (PPI) network and to filter miRNA-mRNA interactions that are not negatively correlated. CancerNet detects synergistic miRNA pairs based on the functional similarity of the targets of the miRNAs and the proximity of these targets in the protein-protein interaction network. This functional association does not necessarily point to synergism in regulating a single gene, but instead, it refers to a group of genes that are regulated together by the same pair of miRNAs.

Finally, Zhang et al. [65] and Zhang et al. [71] uses gene expression data and apply causal inference methods to find synergistic pairs. In Zhang et al. [65], miRNA knockout is simulated using gene expression data to discover synergistic miRNAs. In a more recent study, Zhang et al. [71] extends this work to incorporate the simulation of multiple knockouts of miRNAs. Zhang et al.’s method [65] relies on the IDA method, Interventional calculus when DAG is Absent. Similarly, miRsyn, [71], makes use of the joint-IDA method to identify the synergistic miRNAs. IDA learns causal structure by the PC-algorithm [60] and measures the effects with Pearl’s do-calculus method [11]. In this method, the variables are assumed to follow a multivariate Gaussian distribution, which might not hold in the case of expression data. IDA also assumes there are no hidden effects on the mRNAs, however, the main regulators of the mRNA are actually the transcription factors. The mirSyn method also includes a filtering step that removes transcripts from the analysis that are not significantly associated with survival. mirSyn filters the miRNAs and mRNA that are significantly different in their survival using a Cox regression model. This limits the applicability as patient survival data are not available in many situations and the participants of the synergistic relations are not necessarily associated with survival. Another limitation is that the two works do not check if the binding sites of the candidate miRNAs are overlapping or not. If they are overlapping, it is unlikely that the miRNAs can bind at the same time.

We cast the problem of finding synergistic miRNAs and their mutual target as a statistical dependence discovery problem. Using matched miRNA and mRNA expression data and considering the expression level of each miRNA and mRNA as a random variable, miRCoop finds a specific three-way statistical interaction among these random variables. Here, the pairs of miRNAs have weak or no statistical dependence with the mRNAs, yet when taken jointly, they are dependent on the mRNA. This case is similar to the example wherein adding sugar to coffee or stirring the coffee alone has weak or no effect on the coffee’s sweetness, but the joint presence of the two has a strong effect ([48]). This specific case has not been the focus of previous work. In testing whether the triplets of RNAs have such relationships, we resort to the kernelized Lancaster three-variable interaction developed by Sejdinovic et al. [48]. This test relies on mapping the probability distributions into a reproducing kernel Hilbert space (RKHS). The effectiveness of embedding probability distributions are demonstrated in different machine learning and inference task (reviewed in [53] and [41]). The key idea of embedding probability distributions is to implicitly map probability distributions into potentially infinite-dimensional feature spaces using kernel functions ([52]). Kernel embedding of the probability distributions avoids the restrictive assumptions on the distributions such as variables to be discrete valued, or assumptions on the relationships among the variables such as the linear relationship between variables ([53]). The kernel trick helps us to deal with small sample sizes for the available expression profiles, and it does not require any distributional assumptions. In addition to the expression profiles, miRCoop makes use of the putative target information among the miRNAs and mRNAs.

We apply miRCoop to kidney cancer using tumor samples from The Cancer Genome Atlas (TCGA) project. This analysis discovers 66 putative triplets. We use several validation strategies to check the existence of a regulatory relationship among the triplets’ participants. Investigating the triplets’ RNAs reveal that most of them are individually associated with kidney cancer, as well. Further more, we observe that some miRNA pairs are encoded very close on the genome hinting a tight regulation at the genome level and supporting a functional link among these miRNAs. To check other coordinated regulatory evidence, we investigate whether a shared transcription factor potentially coordinates the miRNAs and RNAs, and interestingly, we observe that this holds for almost all identified triplets. When we explore the relevance of the identified triplets to kidney cancer, our functional enrichment analysis on the largest connected component of miRNA–miRNA–mRNA network highlights critical processes related to kidney cancer, part of which are key driver genes.

## 2 Materials and methods

miRCoop relies on miRNA–mRNA target predictions and kernel statistical tests on the expression data. The overall pipeline of miRCoop is summarized in Fig 1. The first step curates a list of possible triplets based on miRNA target predictions and their predicted mRNAs. In the second step, the candidate triplets are pruned based on expression data. In the final stage, we test the remaining triplets for the statistical dependence relationships. Below, we detail these three steps. Following the miRCoop descriptions, we provide information on the null model used for triplets and the data collection and pre-processing steps when miRCoop is applied to kidney cancer.

**Figure 1:**
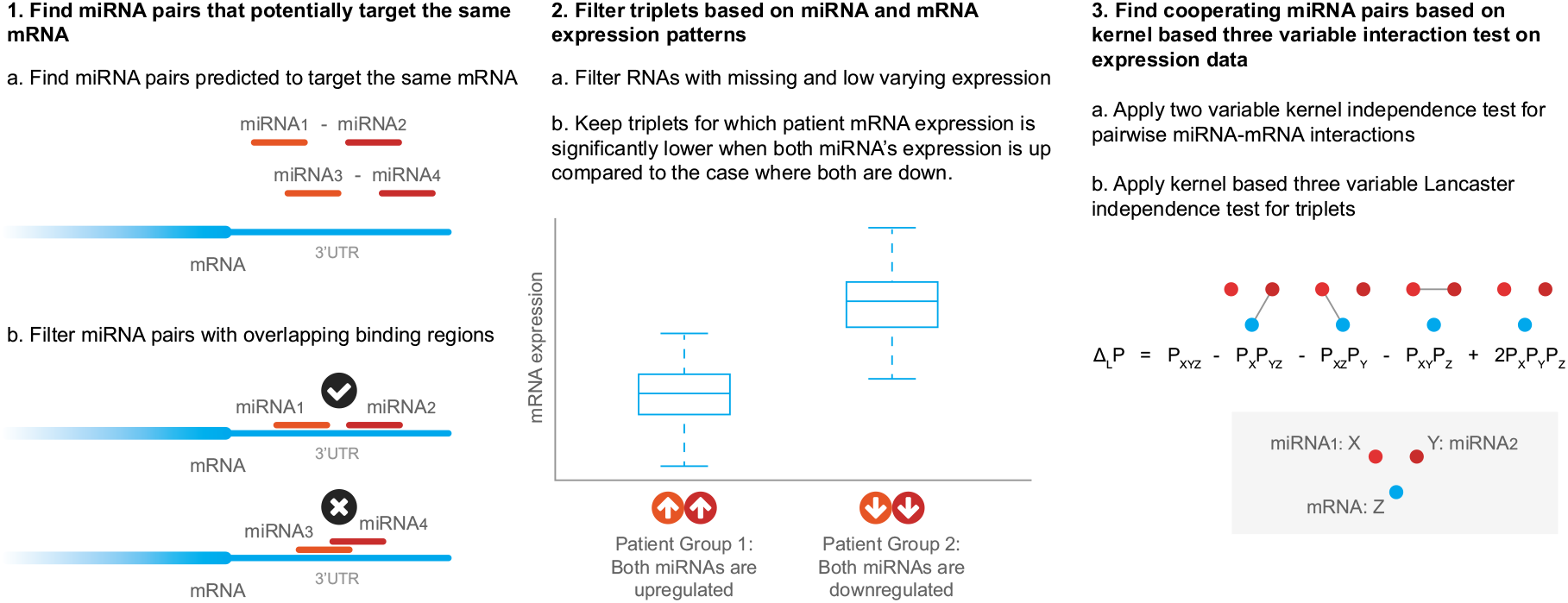
Overview of the methodology. Each box represents a step in the methodology. The first step identifies miRNA pairs that can target the same RNA based on miRNA target predictions. The second step illustrates how the RNAs are filtered based on expression profiles. The third step illustrates the idea of cooperative RNAs. *δL_P_* denotes the Lancaster interaction on the triple of random variables (*X, Y, Z*) is defined as the signed measure. *P_XYZ_* denotes the joint distributions of *X, Y, Z* random variables, while *P_X_, P_Y_* and *P_Z_* are marginal distributions. If the joint distribution P_XYZ_ factorizes into a product of marginals in any way, then *δL_p_* vanishes. This is used as a test statistic to test for independence.

### 2.1 miRCoop Step 1: Identifying Candidate miRNA–mRNA Target Interactions

miRCoop first identifies pairs of miRNA that can bind to a common target mRNA (Fig 1 Step 1). A pair of miRNAs and their mutual target mRNA constitute a candidate triplet. As the interactions of newly cataloged miRNAs are not included in experimentally validated miRNA target interaction databases, we resort to available target prediction algorithms. For this, we use TargetScan ([1]) and miRanda ([7]). When using TargetScan, the 6–mer sites’ predictions are filtered since they are classified as poorly conserved ([1]). Similarly, when running miRanda, to reduce false-positive predictions, we apply stringent parameter settings (a pairing score of > 155 and an energy score of < −20). We consider only the miRNA-mRNA pairs that are predicted by both algorithms. The triplets where miRNA pairs have an overlapping binding site on the target mRNA are also excluded.

Although the target information prunes many combinations, at the end of this step, there is still an extensive list of candidate triplets. For example, when we apply to kidney cancer (see Results), there are 31, 184 potentially interacting miRNA-mRNA pairs, and there are 163, 550 candidate miRNA-miRNA-mRNA triplets.

### 2.2 miRCoop Step 2: Expression Filter

We further narrow down the candidate list based on the expression profiles of the miRNAs and mRNAs measured over a set of samples, Fig 1 (Step 2). miRNAs repress the expression of their target mRNA genes. Therefore, we expect that if both miRNAs are upregulated, the mRNA expression will be lower compared to the case where both miRNAs are down-regulated. To filter for triplets where this expression pattern holds, we divide the samples into different expression subgroups based on the expression states of the potential synergistic miRNA pairs. The first group comprises the samples where both miRNAs are upregulated, and the second group is the subset of samples where both miRNAs are downregulated. In this analysis, miRNA expression in a sample is considered upregulated if it is in the fourth quartile of all samples, and it is deemed to be downregulated if it is in the first quartile. Having grouped the samples, for every triplet in the candidate list, we check whether the mRNA expression levels are significantly lower than the mRNA expression levels compared to the second group. To test for significance, we apply the one-sided Wilcoxon rank-sum test ([56]). This test is used to determine whether two groups of expression values are selected from populations having the same mRNA expression distribution. We apply this test to all possible triplets formed at the end of Step 1 and retain only those for which the null hypothesis is rejected at 0.05 significance level.

### 2.3 miRCoop Step 3: Testing Statistical Interactions of RNAs with Kernel-Based Interaction Tests

For every triplet candidate that passed through Step 2, we perform statistical interaction tests on RNA expression data to find putative miRCoop triplets, Fig 1 (Step 3). We are looking for a high order interaction between the three RNA entities that is while each miRNA expression levels have weak or no interaction on the mRNA expression, the simultaneous interaction of the two miRNAs with the mRNA expression levels is strong.

We denote the two miRNAs’ expression levels as two random variables: *X* and *Y* and define the expression level of the mRNA to be tested as a third random variable denoted as *Z*. We are interested in cases where the miRNAs are pairwise independent with mRNA: *X* ╨ *Y* and *Y* ╨ *Z* but form a mutually dependent triplet because of the following holds: (¬((*X, Y*)╨ Z)). Note that this also corresponds to checking for a V-structure, a directed graphical model with two independent parents pointing to a single child, where the miRNAs’ expression levels (*X* and *Y*) are the parent variables, and the child variable is the mRNA expression level (*Z*) ([48]).

To ensure that the miRNAs have weak influence individually, we first test for pairwise independence among *X* and *Z* and *Y* and *Z*. In each of these tests, the null hypothesis that the two variables are independent. We continue with triplets, where the null hypothesis is accepted. On these triplets, we apply three-variable kernel interaction tests proposed by [48] and considers only triplets that reject the null hypothesis. Here, we provide a brief description of these kernel-based tests and later detail on how we use this method for discovering miRCoop triplets.

#### 2.3.1 Kernel Three-Variable Lancaster Interaction Test

Interaction effects occur when the effect of one variable depends on the value of another variable. In this case, the repression effect of the miRNAs on the mRNA depend on the value of the cooperative miRNA’s expression. An interaction measure is used to test for the existing of such a statistical dependency. An interaction measure associated to a multidimensional probability distribution *P* is a signed measure that vanishes whenever *P* can be factorized as a product of its marginal distributions ([35]). When there are two variables, *X* and *Y*, the Lancaster interaction measure is defined as:

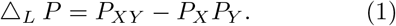

For three random variables, the Lancaster interaction is defined as:

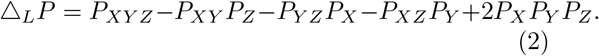

In the above equations, while *P_X_, P_Y_* and *P_Z_* are marginal distributions of the random variables *X, Y* and *Z* respectively. *P_XYZ_* denotes the joint distributions of *X,Y,Z* random variables, *P_XZ_, P_XZ_* and *P_YZ_* are pairwise marginal distributions. The Δ*_L_P* is zero whenever *P* can be factorized as a product of its marginal distributions. A statistical dependency test can be constructed where the Lancaster interaction is the test statistic ([35, 48]). The null hypothesis, in this case, is that the Lancaster interaction is zero. The null distribution of the test statistic can be estimated using a permutation-based approach where the indices of the variable(s) of interest are shuffled ([48]).

Sejdinovic et al. [48] formulate a kernel-based three-variable Lancaster interaction test by embedding the probability distributions into reproducing kernel Hilbert spaces (RHKS). In the general formulation of [48], the three variables to be tested form a random vector of triplets (*X, Y, Z*) ∈ 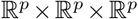, where *p* is the dimension. In our case *p* = 1 as for each miRNA and mRNA there is only one measurement per sample.

The key idea of embedding probability distributions is to implicitly map probability distributions into potentially infinite dimensional feature spaces using the positive definite *kernel functions* [41]. The positive definiteness of the kernel function guarantees the existence of a dot product space H and a feature mapping 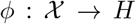 such that *k*(*x_i_, x_j_*) = 〈*ϕ*(*x_i_*), *ϕ*(*x_j_*)〉*_H_*. This dot product can be calculated implicitly with the kernel function without mapping *ϕ* explicitly. This is commonly referred as the *kernel trick* [45]. The mean embedding of a random variable *X* is defined as: 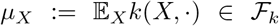. In [48], given observations *X_i_*, an estimate of the mean embedding is 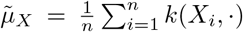. Given an *i.i.d*. sample 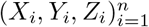, the norm of the mean embedding *μ_L_* is empirically estimated using empirically centered kernel matrices. The empirically centered kernel matrix 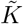 for the kernel *k* is given by:

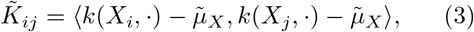

Sejdinovic et al. [48] arrives at the following test statistics for the two variable and three-variable dependency test using the kernel trick:

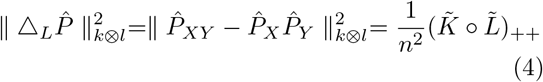

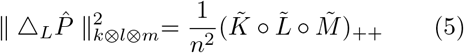

Here, *k* denotes the kernel function defined in domain 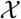 that maps *P_X_* to *H*. Similarly, *l* and *m* are kernel functions defined in domains of 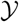 and 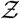, respectively. Using the kernel trick, computations on the probability distributions can be calculated implicitly via kernel matrix operations. The kernel matrix for the kernel *k* is defined as the *n* × *n* matrix *K_ij_* = *k*(*X_i_, X_j_*) computed over all sample pairs, *i,j*. ∘ denotes an Hadamard (elementwise) product of matrices. ⊗ indicates the Kronecker product. **1** denotes an *n* × 1 vector and 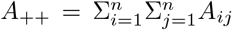 for a matrix *A*. 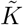 denotes the corresponding centered matrix of *K*, where 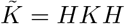 and 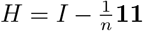 and *I* is the identity matrix.

The same rules apply to the other gram matrices, *L* and *M*. Given an i.i.d. sample 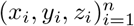, the centered kernel matrices can be computed over the samples. Using equations 5, the norm of the embeddings can be computed using these centered matrices. We refer readers to [48] for the derivation of these equations.

#### 2.3.2 Applying Kernel-based Interaction Tests

We form a random vector of triplets (*X, Y, Z*) ∈ 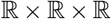. In the pairwise interaction tests, where we test for independence of miRNA expression levels with the mRNA (*X-Z* and *Y-Z*), we consider triplets where the pairwise independence test fails to be rejected (*α* = 0.01). For the three-variable independence interaction test, we only consider, triplets, where the null hypothesis of the three variable interaction is rejected (*α* = 0.01). We use the Gaussian kernel as defined:

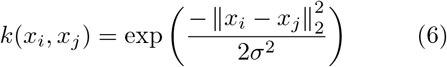
where *σ* is a non-negative real number that determines the width of the kernel. We set *σ*using the median heuristic ([19]). In both tests, the null distribution is estimated by repeatedly permuting the indices of the mRNA expressions of the samples (Z variable), where we set the number of permutations to 2, 000.

### 2.4 A Regression-Based Method

We use a simple alternative strategy to detect synergistic miRNAs using linear regression, which formes a baseline. Starting from the candidate triplets after Step 2 in the miRCoop pipeline, we build three regression models. In the first two models, we model the relationship of miRNAs with the mRNA separately by fitting a linear equation (Eq 7 and Eq 8), where *X* indicates the first miRNA’s expression level and *Y* indicates the second one’s expression. In the third model, in addition to the main effects of the individual miRNAs, we include an interaction term (*X* * *Y*) to measure synergy. A significant, negative, and large in magnitude regression coefficient will indicate a synergy. These three models are given below:

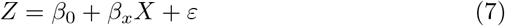

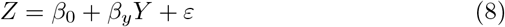

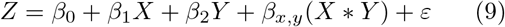

In the fist two models, a significant, negative, and small in magnitude coefficient suggests a weak or non-existing linear relationship between the miRNA and the mRNA. To decide whether the magnitude of a coefficient is large or small, we need to set a cut-off value. Looking at the distribution of the coefficient values of all miRNAs (Supplementary File 1 Fig S5), we pick 0.25. We expect to see a negative correlation between the interacting miRNA-mRNA. Moreover, since we are interested in cases where the miRNA itself shows weak interaction, we keep the triplets for which both miRNA’s regression coefficients are small. Thus, we only consider cases where −0.25 < *β_x_* < 0 and −0.25 < *β_y_* < in the first two models (Equations 7 and 8). Having filtered out triplets for which the above conditions on *β_x_* and *β_y_* does not hold, we fit the third model (Equation 9). For this, we look for cases where *β_x,y_* in Eq. 9 is negative and large. In all cases, we only considered triplets with significant regression coefficients (*p*-value < 0.05). The *p*-value is for the t-statistic of the hypothesis test with the null hypothesis that the corresponding coefficient is equal to zero against the alternative hypothesis that it is non-zero.

### 2.5 The Null Model for miRCoop Triplets

Having identified the triplets, to check the validity of different comparisons with the biological data, we use a random sampling-based approach to obtain a random set of triplets that constitutes the null model. We use these randomized triplets to obtain an empirical null distribution of the test statistic of interest in each case. We construct the randomized triplets as follows: let *S* = *t*_1_, *t*_2_,…, *t_n_* be the set of miRCoop triplets identified. Let the *i*-th triplet member of S denoted by *t_i_* = (*x_i_, y_i_, z_i_*), which include the miRNA pairs, *x_i_, y_i_* and the mRNA z_i_. We sample a collection of *B* random triplet sets of each with size *m*. Let 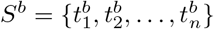 denote the *b*-th random triplet set. For the *b*-th sampling, 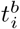 is generated as follows: the mRNA *z_i_* is retrieved, and one miRNA pair of the triplet (*x_i_* or *y_i_*) is selected at random and retained, the paired miRNA triplet is though selected uniformly at random from the set of miRNAs that can potentially target *z_i_, R_z_*, as identified at the end of Step 2 of the miRCoop pipeline excluding the miRNA triplets (Fig 1). By repeating this procedure for all samplings ∀*b* ∈ {1,…, *B*}, we obtain a collection of *B* many triplet sets, 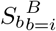 the random counterparts of *S*. We must note that this procedure constructs a rather stringent null model. Note that if we were to select the mRNAs and miRNAs all at random, the miRNAs and mRNAs in these random triplets would likely be unrelated to each other. With such a null model, attaining significance would be essentially trivial.

We calculate the *p*-value of the observed statistic empirically based on this null distribution. To asses the statistical significance of the statistic of interest (such as the number of TFs (See sections 3.2.2, 3.2.1, 3.3)), we obtain an empirical *p*-value by calculating the fraction of the cases in *B* random samplings where the test statistic is as extreme or more extreme than the observed statistic. The null model is constructed with 1, 000 samplings.

### 2.6 Data Collection and Processing

We apply miRCoop on the patient-derived kidney cancer samples. We retrieve the precomputed mRNA and miRNA expression levels from GDC Data Portal on Sept 2^nd^ 2017. mRNAs were quantified using RNA-seq technology while miRNA’s are using miRNA-seq technology. The details of miRNA transcriptomic profiling are described in [12]. The details of data processing can be accessed at https://docs.gdc.cancer.gov/. We utilize the available Reads per million mapped reads (RPM) values and Fragments per Kilobase of transcript per Million mapped reads upper quartile normalization (FPKM-UQ) for miRNA and mRNA expression data, respectively. We only use matched primary tumor solid samples that concurrently include mRNA and miRNA expression data, which contain 995 tumor patient samples. To filter out the non-coding RNA genes from the RNASeq expression data and create an mRNA list, we used GENCODE v27 annotations ([24]).

We eliminate mRNAs with 3’UTRs shorter than 12 nucleotides since two miRNAs cannot bind on such a short sequence at the same time. 3’UTR sequences of mRNAs are obtained using ENSEMBL BioMart ([15]). Genes with expression values lower than 0.05 in more than 20% of the samples are filtered. Expression values are added with a constant 0.25 to deal with the 0 gene expression values, and data are log 2 transformed. We eliminate RNAs that do not vary across samples. For this purpose, we filter RNAs with a median absolute deviation (MAD) below 0.5. In the final set, we have 12,113 mRNAs and 349 miRNAs in kidney tumor samples.

## 3 Results

### 3.1 Overview of the synergistic miRNAs and their targets in Kidney Cancer

We applied the methodology summarized in Fig 1 to find synergistic miRNAs and their targeting mRNAs. The number of candidate triplets that remain after applying each of the filtering steps is provided in Supplementary Fig S1. The target prediction step produces 31, 184 potentially interacting miRNA-mRNA pairs and 163, 550 miRNA-miRNA-mRNA interaction combinations. Following the remaining steps, we arrive at potential 66 synergistic triplets, which also contains 66 different synergistic miRNA pairs. The triplets include 75 miRNAs. The summary statistics of the expression values of these miRNAs are provided in Supplementary File 4. These triplets include 41 different mRNAs, 27 of which are reported to be cancer driver genes for kidney cancer as reported in IntOGen ([22]). The complete list of the identified triplets is listed in Supplementary File 2 and the network for the potential miRNA-miRNA pairs is provided in Fig 2.

**Figure 2:**
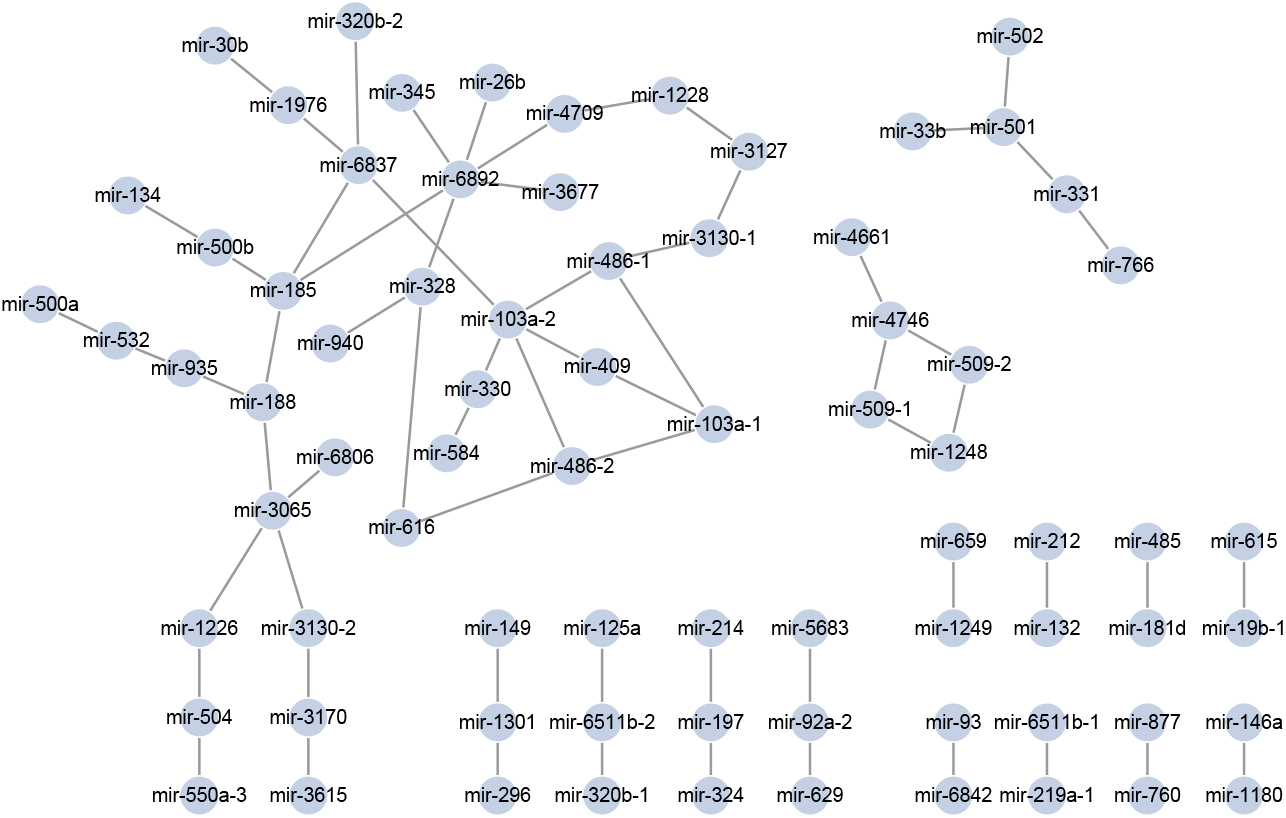
The network shows the miRNA synergistic pairs discovered for kidney cancer. Nodes are miRNAs and edges exist between them if they are participate in at least one miRCoop triplet.

### 3.2 Analysis of the Identified Triplets

There is no available experimentally identified set of synergistic miRNAs to validate the triplets we identified. Thus, we devise several strategies to investigate the miRCoop triplets in light of external biological information. To check the statistical significance of each of these findings, we use a null model by generating random synergistic triplets, which we detail in Methods Section 2.4. Below we describe the biological evidence relevant to the identified triplet set.

#### 3.2.1 Triplets Coded Close on the Genome

Recent studies reveal that miRNAs, which regulate similar processes, are located nearby on the genome ([64]). Others report that miRNAs that are proximally encoded are generally co-expressed, [6], and this indicates a functional link. Thus, we investigate whether the identified synergistic pairs areclose on the genome. Specifically, 10 kb of the transcription start and end sites within the genome are considered spatially close. We retrieve known transcription start and end sites of the miRNA from the Ensembl using biomaRt package [15]. We observe that three of such pairs are very close, and these pairs form two different clusters (Table 1). We find this number is not significantly high (*p*-value = 0.163). The reason could be that most reported since the synergistic miRNAs are not necessarily acting *cis*. However, those that we observe close in the genome are interesting. These clustered miRNAs are part of the miR–500 and miR-132 miRNA family. There is evidence that miR-500 family takes part in kidney cancer ([39, 51, 17]). Genomic locations of these synergistic miR-500 family miRNAs are illustrated in Fig 3.

**Figure 3:**
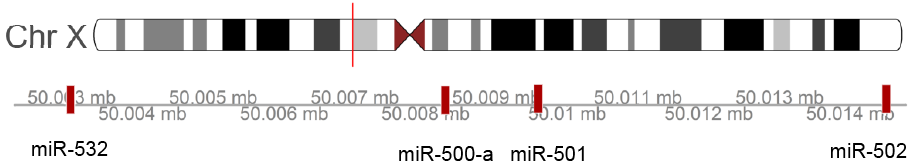
The red marks on the genome axis indicate the miRNA positions of the synergistic miRNAs that belong to the miR-500 family. The figure was created with Gviz package in R ([23]).

**Table 1:** The list of identified synergistic miRNA pairs in kidney cancer that are encoded close in the genome and their family information.

| miRNA | miRNA | Distance | miRNA Family |
| --- | --- | --- | --- |
| miR-132 | miR-212 | 262 | miR-132 |
| miR-501 | miR-502 | 4792 | miR-500 |
| miR-500a | miR-532 | 5192 | miR-500 |

#### 3.2.2 Common TF Regulating the RNAs of the Triplets

Coordinated regulation of genes by a shared transcription factor often indicates a functional link among the genes. To this end, we checked if the identified triplets were potentially co-regulated by a common TF. To be able to conduct this analysis, we first curated a set of regulatory interaction for TFs regulating miRNA and mRNA.

Since none of the TF-miRNA databases are comprehensive, we use two strategies to curate TFs regulating miRNAs. We first use data from available databases; this includes TransmiR ([29]), PuTmiR ([28]) and miRTrans ([27]) databases. miRTrans database provides a list of TFs-miRNA interactions for different tissues and cell types; we only consider the renal cell lines HRE, human Renal Epithelial Cells, and RPTEC, hTERT immortalized human renal proximal tubular epithelial cell lines. Secondly, we compile the list by analyzing ChIP-Seq data made available by the ENCODE project ([26]) and TF motifs. We retrieve the transcription start sites from Ensembl employing the biomaRt package[15] and 10,000 upstream regions of transcription start of the gene in question using the R package GenomicRanges ([66]). We check if this region overlaps with the ChIP-Seq peaks in three kidney-related cell lines (HRE, HEK293B, HEK293) using the GLANET tool [70]. We compile the TF-miRNA interactions from the databases and discovered through analyzing ChIP-Seq data. For the list of TFs regulating mRNAs, we conducted the same analysis on the ChIP-Seq data. Furthermore, we scan the promoter regions with the TFs’ binding motifs with JASPAR ([67]) and Hocomoco v.11 ([68]) using the FIMO ([69]) tool with default parameter settings. If the TF has a reported overlapping peak with the promoter and there is a matching motif inside the peak, we consider this as a TF-mRNA interaction.

Finally, we check if there is at least one common TF that regulates all RNAs (the miRNAs and the mRNA) that participate in a triplet interaction. In checking this, we disregard POL2 as it will overlap with promoter regions without any relevance to the functional cooperation. We observe that for 64 of the 66 triplets, all triplets participants are coregulated by at least one common TF. This number was significantly high at test level (*p*-value = 0.04. The list of the TF-triplet relations is provided in Supplementary File 3.

We inspect a subset of the TF-triplet regulations more in detail. Recent studies report that the transcription factors, HIF1A ([30]), TFEB, TFE3 and MITF ([46]) are related to the kidney cancer. We check if there are any miRCoop triplets regulated by these transcription factors. These four TFs are linked to a total of 27 triplets (out of 66) and the sub-network includes triplets that are coordinated by multiple of these TFs Fig 4. Additionally, we observe that the mRNA pairs that are encoded very closely on the genome (see Section 3.2.1) are among them and have shared TFs. For example, TFE3 potentially regulates both miR-501, miR-502, miR-500a, and miR-532. Apart from this, interestingly, TFs related to the Xp11 translocation renal cell carcinoma disease (TFEB, TFE3 and MITF) ([46]) regulate the same triplets: (miR103a.1, miR486.1, PDZD8) and (miR103a.1, miR330, AMOT).

**Figure 4:**
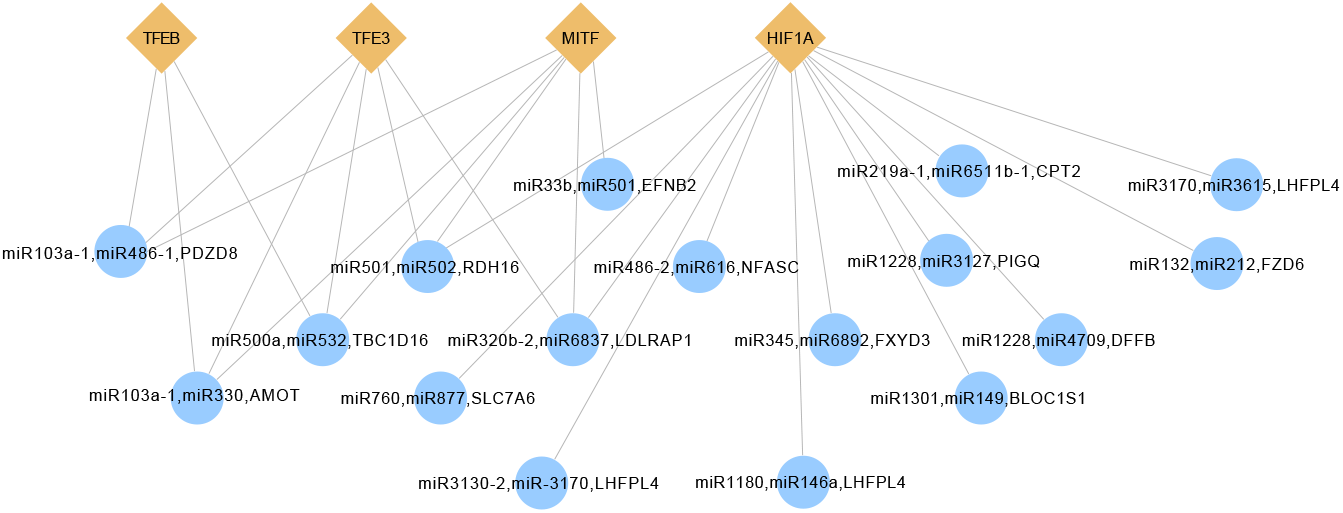
The network for the synergistic triplets regulated by common TFs. Each node denotes a triplet or a TF while an edge represents an interaction between them. TFs are represented by the diamond-shaped yellow symbol, and triplets are represented by the circle-shaped blue symbol.

#### 3.2.3 Functional Enrichment Analysis

Synergistic miRNAs are likely to regulate genes with the same or similar functions ([59]). To investigate the potential biological process related to the synergistic miRNAs, we conduct a functional enrichment analysis on them. To this end, we first build a miRNA-miRNA-mRNA network. In this network, nodes are miRNAs and mRNAs, and an edge exists between two RNAs whenever they are connected through at least one miRCoop triplet. A connected component analysis reveals that the network includes 13 connected components. Interestingly, the largest connected component covers a very large fraction of the all miRNAs: of the 75 miRNAs that is part of the triplets, 41 of them were in this largest connected component (*p*-value = 0.032) (Fig 5 (a)) indicating that the cooperative miRNAs are directly or indirectly linked to each other through their targets.

**Figure 5:**
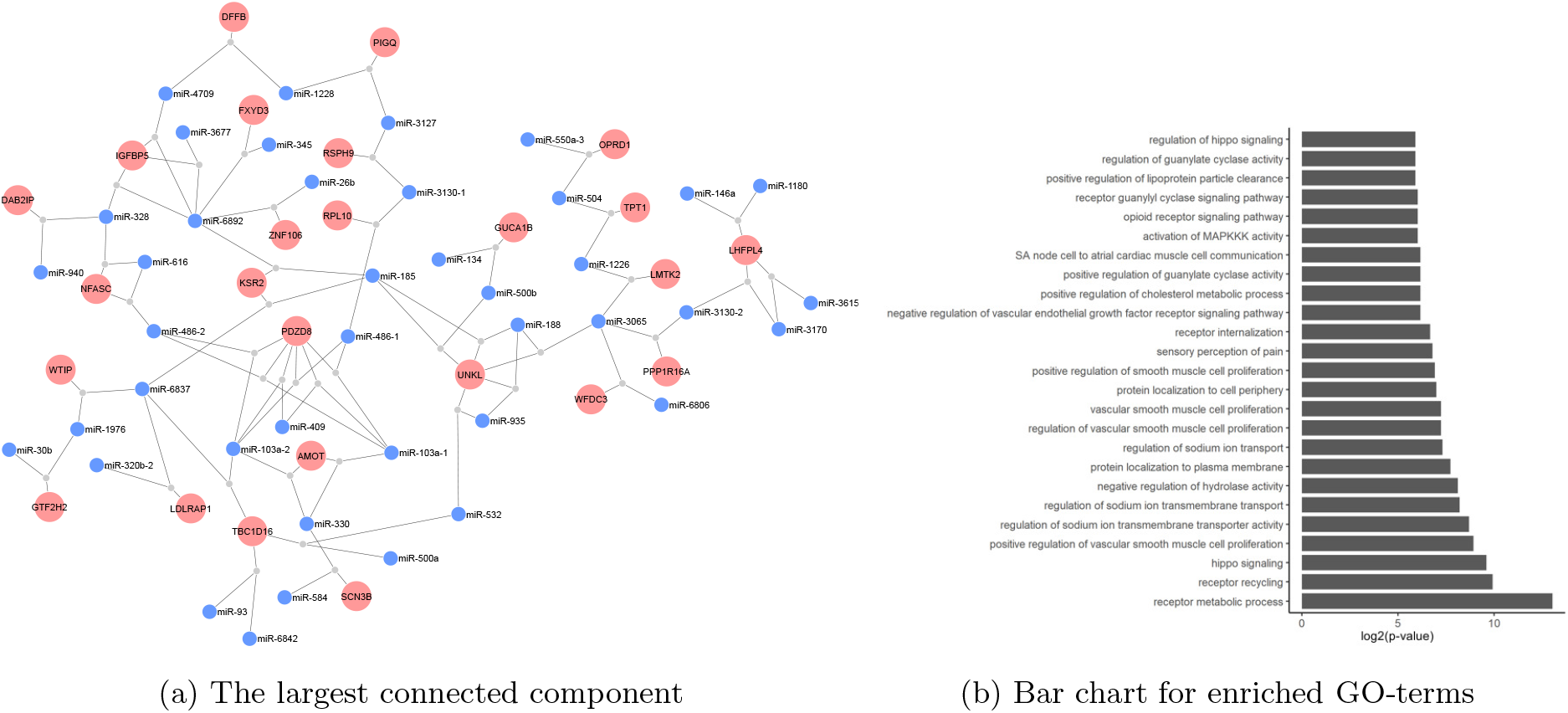
a) The the largest connected component of miRCoop triplet network. Blue nodes are miRNAs and pink colors are mRNAs of the triples. The gray nodes are dummy nodes to connect the triplet miRNAs to the target mRNA. b) Bar chart for the top 25 enriched biological process GO-terms (*p*-value < 0.05) found over the mRNA set of the largest connected component in part a). The x-axis shows log 2 transformed p-values.

Gene ontology (GO) enrichment analysis is GO analysis is conducted with clusterProfiler [63] package in R. We use the default background set that is all human genes annotated by GO Biological Process in the org.Hs.eg.db database [10]. We set a *p*-value cut-off of 0.05, an FDR cutoff 0.05 and used Bonferroni multiple hypothesis test correction.

The GO enrichment analysis reveals that the set of mRNAs are enriched in processes that are associated with kidney cancer. For example, metabolic networks such as the receptor metabolic processes, the negative regulation of hydrolase activity, and the regulation of metal ion transport are all found to be enriched (Fig 5 (b)). Among these, sodium and guanylate related processes are particularly reported as important in kidney cancer ([31, 43, 42]). Also, the genes that are already known as the drivers of renal cancer are annotated with these processes. An example is Angiomotin (AMOT). [37], reports that AMOT is an oncogene in renal cancer. AMOT family members are also reported to promote cancer initiation in other cancers and act as a tumor suppressor in some others ([62, 38]).

Interestingly, the brain related biological processes such as layer formation in the cerebral cortex, synapse organization, embryonic brain development are found to be enriched (Fig 5(b)). It is reported that brain metastasis is common for renal cancer and in 15 % of cases, the tumor metastasizes to the brain ([58]). Additionally, brain metastasis is the main cause of morbidity and mortality in kidney cancer ([58]). These findings suggest that mRNAs regulated by the synergistic triplets may participate in the mechanisms that promote brain metastasis.

### 3.3 Comparison with Other Methods

We first check the overlap of the identified triplets with with that of other computational methods available in the literature: CancerNet ([40]), TriplexRNA ([47]) and miRsyn [71] and a simple regression-based model (See Section 2.4). CancerNet provides cancer-specific pre-computed synergistic pairs of miRNAs. Of the 66 synergistic miRNA pairs, miRCoop identifies, 35 of them are also suggested by CancerNet. This overlap is highly significant based on the sampling-based null model with a sampling size of 1, 000 (*p*-value ≤ 0.001). We also check the overlap with the synergistic pairs provided by TriplexRNA; however, we do not find a significant overlap. One reason for this minimal overlap could be that TriplexRNA triplets are not cancer-specific while the triplets we identified are. mirCoop do not have any overlapping triplets with the miRsyn and the regression-based model. The sizes of the overlaps are summarized in the Venn diagram in Fig 6.

**Figure 6:**
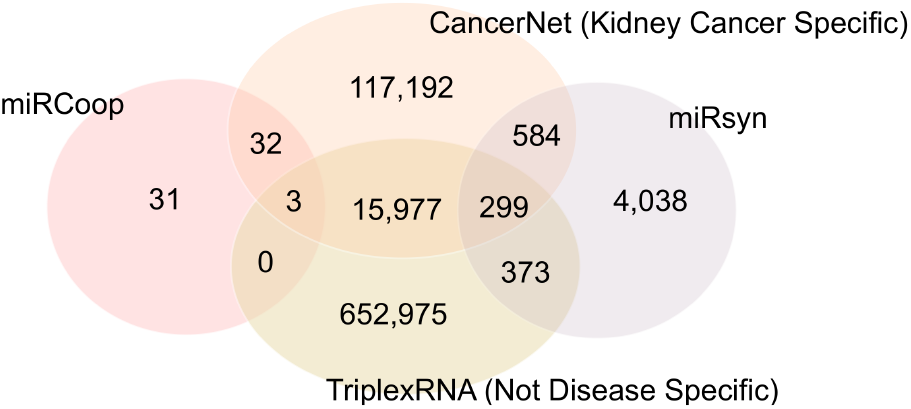
The Venn diagram for the number of synergistic miRNAs that are detected by different methods. In the TriplexRNA database, we utilize TEC predictions.For Cancernet predictions, kidney cancer specific results are used. miRsyn results are obtained by running the miRCoop tool on the TCGA kidney data.

We also compare mirCoop with mirSyn [71]. We run miRSyn on the same data using the same data that were described in Section 2.6 using suggested parameters. When considered only the pairwise synergistic pairs, mirSyn return 5,284 miRNA-miRNA-mRNA triplets that involves 1, 079 synergistic miRNAs. There was no overlap of the triplets with mirSyn, although there were 11 miRNAs that overlap. MiRsyn employs a filtering step based on Cox regression, thus eliminates transcripts that do not prognostic value, and the methods used are different. This could be partly the reason why there is no overlap.

We built the regression model (Section 2.4) to constitute a baseline to assess whether mirCoop triplets could be identified with an easier strategy. This simple method results in 24 potential triplets, which do not have overlap with mirCoop triplets or any of the tools we compare the miRCoop against (shown in the Venn diagram provided in Supplementary File 1 Fig S2).

We also compare mirCoop with mirSyn and the regression method in terms of the genome and the TF relations. For the regression model, none of the synergistic miRNAs are located close by on the genome. We also find none of the triplets are regulated with a common transcription factor. Thus, we conclude that this simple strategy is not sufficient to solve this problem. This method has various limitations, as well. It assumes that the nature of the statistical interaction of the individual miRNAs and mRNAs are linear. Also, the linear regression model assumes that the mRNA expression values are normally distributed (as the errors are assumed to be normally distributed in the standard regression). We also need to set arbitrary cutoffs on the coefficients to decide the strong and weak interactions.

We analyzed the synergistic miRNAs in miRsyn that are encoded close-by on the genome. We observe that 10 out of 1, 079 synergistic miRNA pairs suggested by miRsyn [71] are close-by on the genome. The number of 10 was not significantly high at confidence level 0.05 when tested with the permutation test. Next, we check if miRsyn triplets are co-regulated by common TFs. We conducted a statistical test exactly as we did in miRCoop to check if this number is interesting hight. The synthetic interactions in the null model are constructed as described in section 2.5. The test fail to reject the null hypothesis with *P* – value = 1.00. On average, (on average 3, 810 triplets are co-regulated by at least one common TF) calculated over 1000 samplings.

## 4 Conclusion

Post-transcriptional regulation of gene expression mediated by miRNAs is a fundamental mechanism to fine-tune the protein levels in the cell. The dysregulation of miRNA expression is also associated with many diseases, including cancer. miRNAs are not only potential biomarkers, but are also investigated as promising therapeutic agents; therefore, understanding the intricate ways they work is essential for both expanding the knowledge on how cells work and translational purposes. In this work, we focus on the synergistic target regulation of miRNAs. To identify synergistic miRNAs pairs and their target mRNA, we propose a novel method: miRCoop. MiRCoop uses kernel-based statistical interactions and finds triplets where miRNA expression levels have no or only weak statistical dependencies with the mRNA levels but are jointly dependent. miRCoop makes use of both RNA sequence information for target predictions. We demonstrate the use of miRCoop in kidney cancer and find triplets with supporting biological evidence on their relevance. The work presented here can be extended in several directions.

Although it is possible to extend the Lancaster interaction test to detect interactions of higher-order, where more than two miRNAs regulate a mutual target, it would necessitate significantly higher computational resources. Therefore, at the moment, we limit our analysis to the three-variable interaction case. Finding higher-order interactions with ways to reduce the computation time can be a future line of research.

In this work, we focus on miRNAs that regulate a target gene directly which is those that simultaneously bind to the target. There are other ways in which pairs of miRNAs regulate a common target. For example, while one miRNA targets the gene of interest directly, another miRNA can target the transcription factor that regulates this gene ([9, 65]). In this case, one miRNA is acting directly on the target, and the partner miRNA is acting indirectly through the transcription factor. An additional line of future work would be to extend miRCoop to discover these multi-level regulations.

Since there are no synergistic interaction datasets that are readily available, we devise various strategies to check if the identified triplets and the individual RNAs are implicated in kidney cancer and have any biological evidence supporting a functional theme or regulatory interaction among themselves. When experimentally validated synergistic miRNA pairs become available, the analysis can be further improved in the light of the ground truth interactions. For example, we set the kernel width using the median heuristic, but the width could be tuned when an extensive set of ground truth interactions become available.

In the present study, to establish target relations between the miRNA and the mRNAs, we use two target prediction algorithms. We resort to prediction algorithms because newly cataloged miRNAs – which is valid for the TCGA profile miRNAs - are not incorporated in the experimentally validated miRNA target interaction datasets. To reduce the number of false positives, we only consider target interactions that are supported by both algorithms concurrently. The availability of experimentally validated miRNA-mRNA interactions should strengthen the outlined methodology. The number of synergistic triplets can be adjusted by changing the significance levels applied in the statistical tests. For example, in a scenario where a large number of interactions can be verified experimentally at scale, these settings can be adjusted to output more triplets.

In this study, we use all kidney cancer patients to have a large sample size. However, the set of synergistic miRNAs could be different for different subpopulations of the same cancer type. As more data accumulate, the analysis can be repeated over subtypes of cancer.

Although we showcase results for kidney cancer, the method can be applied to other diseases as well as healthy cells. The method can be extended to work with other species other than human whenever the expression profiles of matched miRNAs and mRNAs are available for over a set of samples, and miRNA target predictions are available.

## Acknowledgments

The results shown here are in whole or part based upon data generated by the TCGA Research Network: https://www.cancer.gov/tcga. We thank Sabanci University and Bilkent University for internal funding support. This research was supported in part by the Intramural Research Program of the NIH, NCI, Cancer Data Science Lab.

