## Supplementary File 1 for "miRCoop: Identifying Cooperating miRNAs via Kernel Based Interaction Tests"

### Supplementary Materials for ‘miRCoop : Identifying Cooperating Pairwise miRNAs via Kernel Based Interaction Test ’

Gulden Olgun[1], Oznur Tastan[2*]
[1] Cancer Data Science Lab, National Cancer Institute, NIH, Bethesda, MD, 20892
[2] Faculty of Engineering and Natural Sciences, Sabanci University, Tuzla, Istanbul, Turkey
*

Table S1: Details of the data set that are utilized in the analysis.

| Primary Site | Data type | Date of download | Data portal | Webpage |
|---|---|---|---|---|
| Kidney | mRNA Expression | 2.09.2017 | TCGA v2 | https://portal.gdc.cancer.gov/ |
| Kidney | miRNA Expression | 2.09.2017 | TCGA v2 | https://portal.gdc.cancer.gov/ |
| Kidney | Clinical Data | 2.09.2017 | TCGA v2 | https://portal.gdc.cancer.gov/ |

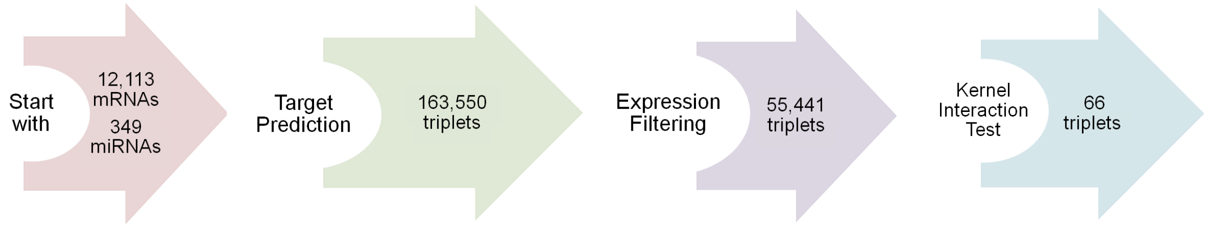


Figure S1: The number of remained triplets after each step in the pipeline. Also, the number of available RNAs that participate in the analysis is provided.

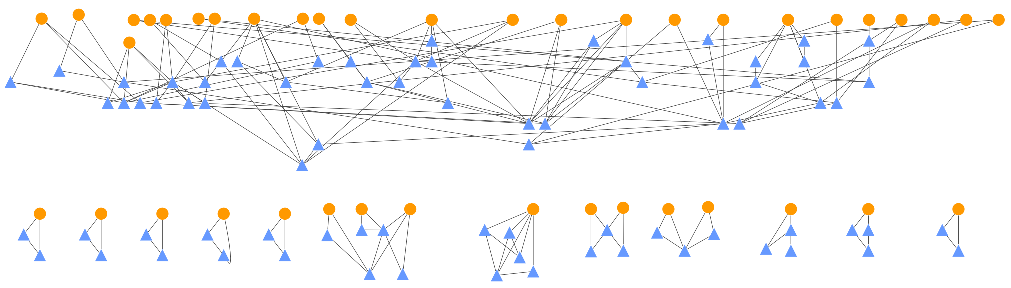

Figure S2: The connected components on the miRCoop triplets. The orange circle shaped nodes represent the mRNAs and the blue triangle shaped ones represent the miRNAs. Edges exist if they are part of at least one triplet.

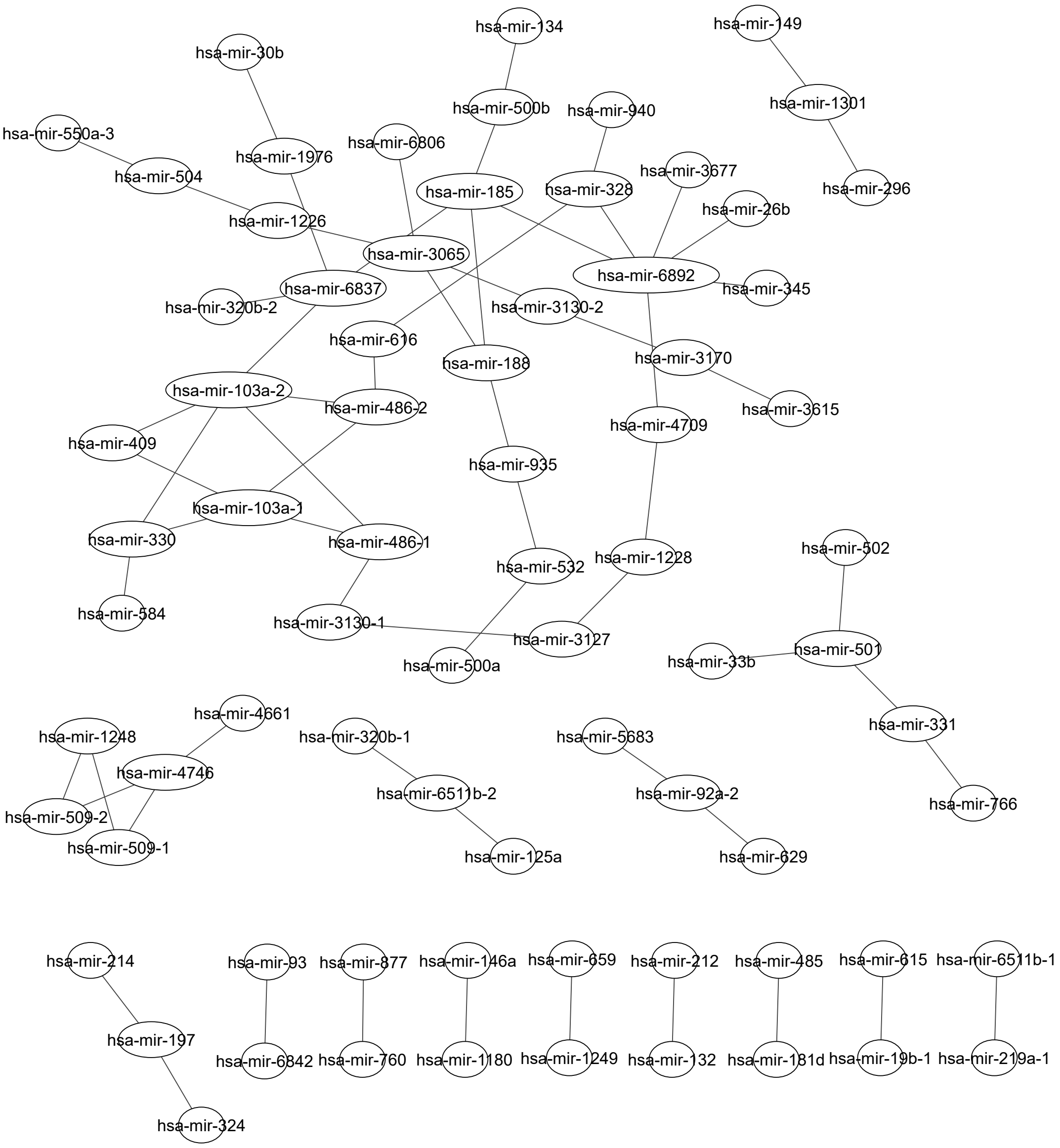


Figure S3: The network of all identified synergistic miRNA pairs. Each node is an miRNA and edges exist if they are predicted to be in a synergistic relationship in kidney cancer patients.

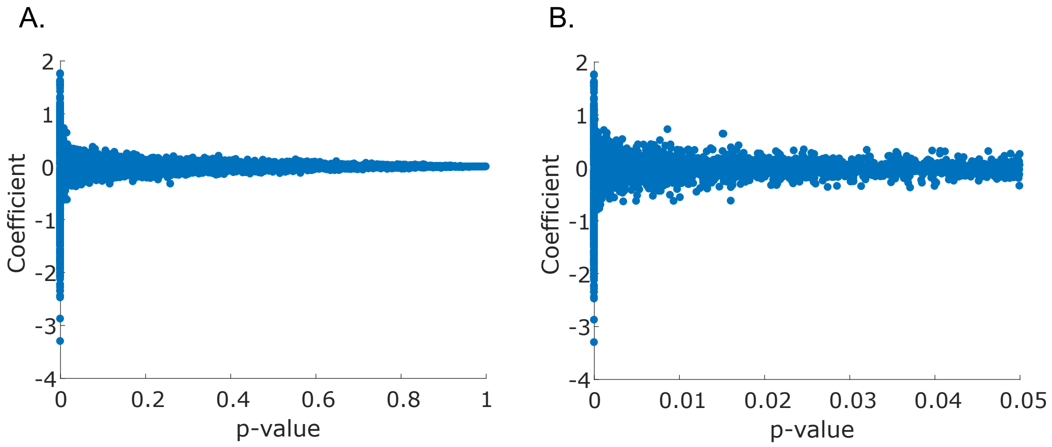


Figure S4: The distribution of the coefficient, $\beta$, for miRNA-mRNA repression models. (A) presents all coefficient values, $\beta$, for each miRNA-mRNA interactions whereas (B) shows only the statistical significant (t-test, $p$-value $< 0.05$) coefficients.
